# Archaeological preservation of amelogenesis pathways

**DOI:** 10.64898/2026.03.25.713862

**Authors:** Ragnheiður Diljá Ásmundsdóttir, Gaudry Troché, Jesper V. Olsen, Marina Martínez de Pinillos, María Martinón-Torres, Sarah Schrader, Frido Welker

## Abstract

Dental enamel, the hardest mineralised tissue in the human body, has proven to be an excellent source of ancient proteins, which have been found to survive within dental enamel for at least twenty million years. In archaeological and palaeontological contexts, the enamel proteome is generally considered to be rather small, consisting of about twelve proteins, most of which are unique to enamel. During amelogenesis these proteins undergo *in vivo* digestion by matrix metalloproteinase 20 (MMP20) and kallikrein 4 (KLK4) as well as serine phosphorylation by ‘family with sequence similarity member 20-C’ (FAM20C) that alter their characteristics. Gaining knowledge of the previously understudied influence of amelogenesis on the archaeological human dental enamel proteome could benefit various palaeoproteomic analysis, especially in an human evolutionary context. Here we present archaeological dental enamel proteomes and explore protein cleavage patterns and sequence coverage to estimate the effects of *in vivo* digestion, as well as explore phosphorylation patterns. Additionally, we present a new marker based on phosphorylation to estimate genetic sex.

## 1. Introduction

Dental enamel is the hardest mineralised tissue in the human body, only composed of 1-2% organic material (Castiblanco et al., 2015). Humans are diphyodont, losing their first set of dentition through an exfoliation process where the roots are reabsorbed, leaving only the enamel crown and occasionally a small portion of the dentine and/or enamel-dentine junction (EDJ) underneath, before acquiring the second set, the permanent teeth (Sahara et al., 1993). Mature dental enamel is acellular, void of DNA, and rich in highly unique proteins (Demarchi et al., 2016). The enamel has been widely used in human evolutionary research, mainly through its morphology and more recently palaeoproteomics (e.g. (Madupe et al., 2025; Tsutaya et al., 2025; Welker et al., 2020)), as the extracted ancient enamel proteomes are shown to contain a few phylogenetically informative sites identified both from mass spectrometry-based proteomic data as well as genomic translations (Madupe et al., 2025; Tsutaya et al., 2025; Zanolli et al., 2017). Dental enamel has proven to be an excellent source of proteins from ancient material, with the oldest hominin proteins identified from ca. 2 million year old enamel sample from *Paranthropus robustus* (Madupe et al., 2025) and the oldest faunal enamel proteins at over 20 million years old (Paterson et al., 2025). Within the palaeoproteomic field, dental enamel is further gaining popularity for its use in estimating the genetic sex of archaeological remains (Blacka et al., 2025; Cintas-Peña et al., 2023; Koenig et al., 2024; Kotli et al., 2024; Lugli et al., 2019; Madupe et al., 2025; Mikšík et al., 2023; Parker et al., 2019; Rey-Iglesia et al., 2025; Stewart et al., 2016, 2017; Tsutaya et al., 2025; Welker et al., 2020; Ziganshin et al., 2020). This is possible as the dental enamel contains two amelogenin proteins, one that is located on the X chromosome (AMELX) and one that is located on the Y chromosome (AMELY) (Santos & Line, 2006).

The enamel proteome is generally considered to be rather small and composed of highly unique proteins not expressed elsewhere in the body. During enamel formation, amelogenesis, five proteins are secreted by ameloblasts, the main cell type in enamel. These proteins are amelogenin (both AMELX and AMELY), ameloblastin (AMBN), amelotin (AMTN), and enamelin (ENAM) (Kawasaki & Weiss, 2003). During amelogenesis these proteins undergo serine (S) phosphorylation by the influence of the protein ‘family with sequence similarity member 20-C’ (FAM20C), which enhances their mineral binding affinities (Dong et al., 2022; Ma et al., 2016; Tagliabracci et al., 2012). FAM20C has not been reported in deep-time palaeoproteomic studies. The ameloblast secreted proteins also undergo *in vivo* digestion by two proteases, matrix metalloproteinase 20 (MMP20; also known as enamelysin) and kallikrein 4 (KLK4), that are unique to the dental enamel. These two proteases cleave the proteins during the final stages of amelogenesis, first with MMP20, which subsequently activates KLK4 (Simmer & Hu, 2002). Peptides belonging to these proteases are also reported in enamel proteomes, with MMP20 identified more often due to its higher concentration compared to KLK4 (Simmer & Hu, 2002; Yamakoshi et al., 2013). How these two proteases work together, and what their exact cleavage sites are, have been explored in both experimental and computational settings (Chun et al., 2010; Iwata et al., 2007; Nagano et al., 2009; Vetyskova et al., 2024; Welker et al., 2020; Yamakoshi et al., 2006), however, it is not fully known whether these cleavage patterns preserve in archaeological proteomes.

Here we present the recovered human enamel proteome from ten archaeological adult permanent maxillary incisors, as well as ten modern deciduous maxillary incisors. We observe a larger number of peptides within the permanent dental enamel. Further research is, however, needed to determine the potential differences between deciduous and permanent tooth proteomes, as the differences observed here cannot be fully explained due to differences in the chronological age of the groups of teeth. From the adult archaeological dental enamel we observed patterns related to amelogenesis, related to protein cleavage patterns and amino acid coverage, similar to experimental porcine data (Chun et al., 2010; Iwata et al., 2007; Nagano et al., 2009; Vetyskova et al., 2024; Yamakoshi et al., 2006, 2013), as well as serine phosphorylation. Additionally, we present a marker for genetic sex estimation based on serine phosphorylation.

## 2. Material and methods

### 2.1 Material

Ten deciduous maxillary incisors (five female and five male) as well as ten permanent maxillary incisors (four female and six male) were selected for this study (SI Table 1). All teeth were either left or right maxillary first incisors, with the exception of individual 13 where the left maxillary second incisor was used. All teeth were healthy and had minimal to no signs of caries and other dental diseases.

The deciduous teeth are modern, from the Ratón Pérez Collection, Centro Nacional de Investigación sobre la Evolución Humana (CENIEH), Spain. This collection was established as a reference collection of deciduous teeth for various archaeological and anthropological studies (Martínez de Pinillos et al., 2021). Children throughout Spain have donated their exfoliated teeth to the collection, with parental and/or guardian consent. Information about age at exfoliation and biological sex was collected upon donation. No other personal information was used for this study.

The permanent teeth are archaeological, originating from the graveyard of the Saint Eusebius church in Arnhem, the Netherlands, which was in use during the Medieval and post-Medieval periods (1350-1829 AD) (Zielman & Baetsen, 2020). All permanent teeth belong to skeletons with estimated age ranging from 18 to 49 years. Both biological age and biological sex were estimated using standard osteoarchaeological methods (Buikstra & Ubelaker, 1994) prior to this work as described in (Ásmundsdóttir et al., 2025).

### 2.2. Methods

#### 2.2.1. Protein extraction

Enamel samples were acquired by cutting off one corner of either the distal or mesial lobe using a dental drill, close to the incisal surface, and avoiding the dentine. In case of very worn teeth, a section along the labial incisal surface was made. All samples, ranging from 2 to 16 mg, with an average of 5.5 mg (SI Table 1), were placed in microtubes (Protein LoBind

Tubes, Eppendorf). Drilling instruments and table surfaces were cleaned between sampling of different specimens, first with 5% bleach, followed by 70% ethanol. One laboratory blank was added after sampling. The samples, alongside a laboratory blank, were demineralised using 1 mL of 5% HCl (Sigma Aldrich) for 48 hours at 3°C. After 24 hours the acid was removed and stored at −20°C, whereafter a fresh 1 mL of 5% HCl was added. Following demineralisation, 0.5 mL from each fraction was cleaned and desalted using in-house made C18 StageTips (Taurozzi et al. 2024).

#### 2.2.2. LC-MS/MS

Peptides were eluded from StageTips with 30 µL of 40% ACN and 0.1% TFA, and then evaporated to approximately 1 µl before being reconstituted with 20 µL of 0.1% TFA and 5% ACN. Peptides were separated with an EasyLC system (Thermo Fisher Scientific) coupled to an Exploris 480 mass spectrometer (Thermo Fisher Scientific) using an in-house made column, packed with 1.9 µm C18 beads with pores of 120 Å (ReproSil-Pur, C18-AQ, Dr. Maisch).

Two mobile phases were used during the liquid chromatography. Mobile phase A consisted of 0.1% formic acid (FA; LC-MS grade FA, Thermo Fisher Scientific), and mobile phase B consisted of 80% ACN (LC-MS grade ACN, VWR) and 0.1% FA. Injection of 3 µL eluded peptide per sample at a flow rate of 500 nL/min was separated on a 77 minute gradient with increasing concentration of mobile phase B, from 5% to 30% over 50 minutes, from 30% to 45% for 10 minutes, prior to a wash at 80% for a duration of 5 minutes and re-equilibration at 5% for 5 minutes. Mass spectrometry was performed in positive ionisation mode at +2000 V. The ion transfer tube was maintained at 275°C. The first MS scan (MS1) ranged from 350 to 1400 m/z with Orbitrap resolution of 120,000. The top ten ions, in data dependent mode (DDA), with minimal intensity of 20,000 and charge states between two and six were selected for fragmentation. MS/MS (MS2) for the selected ions were generated using higher-energy collision dissociation (HCD) set at 30% with an isolation window at 1.2 m/z and resolution of 60,000.

#### 2.2.3. Data analysis

Mass spectrometry raw data (.raw format) were analysed in three different searches using two types of software. One search was performed using MaxQuant (version 2.1.30) (Cox & Mann, 2008). The database used for the search consisted of protein sequences of the following proteins, as described in (Welker et al., 2020), and their isoforms downloaded from Uniprot.org: ALB (P02768), AMBN (Q9NP70, Q9NP70-2), AMELX (Q99217, Q99217-2, Q99217-3), AMELY (Q99218, Q99218-1), AMTN (Q6UX39, Q6UX39-2), ENAM (Q9NRM1), COL1A1 (P02452), COL1A2 (P08123), COL17A1 (Q9UMD9), Dipeptidyl peptidase 1 (CTSC, P53634), KLK4 (Q9Y5K2, Q9Y5K2-2), MMP20 (O60882), ODAM (A1E959), and TUFT1 (Q9NNX1, Q9NNX1-2, Q9NNX1-3). Deamidation (NQ), oxidation (MP), and phosphorylation (STY) were set as variable modifications. The internal MaxQuant contaminant list was included and the search conducted in unspecific digestion mode. Other settings were left at default.

Two searches were performed using PEAKS 7 (Zhang et al., 2012). The first PEAKS search was performed against the whole Uniprot human reference proteome (UP000005640; downloaded September 5th 2023), canonical sequences only with the exception of isoforms for AMELX and AMELY. The second PEAKS search was performed using a database of all proteins identified in the first PEAKS search and all their isoforms. Both PEAKS searches were performed in “none enzyme” mode with the following variable modifications: deamidation (NQ), oxidation (MP), and phosphorylation (STY). The PEAKS searches included a *de novo* search, peaks search, PTM search and spider search. Other settings were left at default. All results were exported with 1% peptide false discovery rate (FDR).

Data analysis was performed with R (version 4.2.2) (R Core Team, 2022) using RStudio (version 2022.02.4) (RStudio Team, 2022). The following packages were used: *Janitor* (version 2.2.0) (Firke, 2023), *Tidyverse* (version 2.0.0) (Wickham et al., 2019), *ggpubr* (version 0.6.0) (Kassambara, 2023), *Patchwork* (version 1.3.0) (Pedersen, 2024), *Paletteer* (version 1.6.0) (Hvitfeldt, 2021), and *pheatmap* (version 1.0.12) (Kolde, 2019).

Mass spectrometry data (.raw format), as well as MaxQuant and PEAKS 7 output, have been deposited to the ProteomeXchange Consortium via the PRIDE partner repository (Perez-Riverol et al., 2022) with the dataset identified PXD075705.

## 3. Results

For the twenty generated dental enamel proteome extracts we conducted a quality control based on the number of MS/MS acquired and the percentage of MS/MS identified (Figure 1), based on MaxQuant output. We observe full separation of dental enamel extracts and the laboratory blank for both MS/MS acquired and the percentage of MS/MS identification. Near full separation is observed between deciduous tooth and permanent tooth enamel extracts based on the percentage of identified MS/MS, where a singular permanent tooth extract is more similar to the deciduous tooth extracts. The percentage of identified MS/MS of the permanent enamel extracts (17-30%) is comparatively high for palaeoproteomic studies. The deciduous enamel extracts have between 5 and 17% identification rate for MS/MS, which is within the upper range expected for palaeoproteomic data (Chiang et al., 2024), but significantly lower than that observed for the permanent teeth. These deciduous teeth extracts are, however, from modern teeth and the identification percentage can therefore be considered low in comparison to meta-analysis of MS/MS identification success rates reported for modern proteome studies (Chiang et al., 2024). This is most likely due to the absence of a laboratory enzymatic digestion step that causes many peptides and their fragment ion spectra to be unidentifiable.

**Figure 1:**
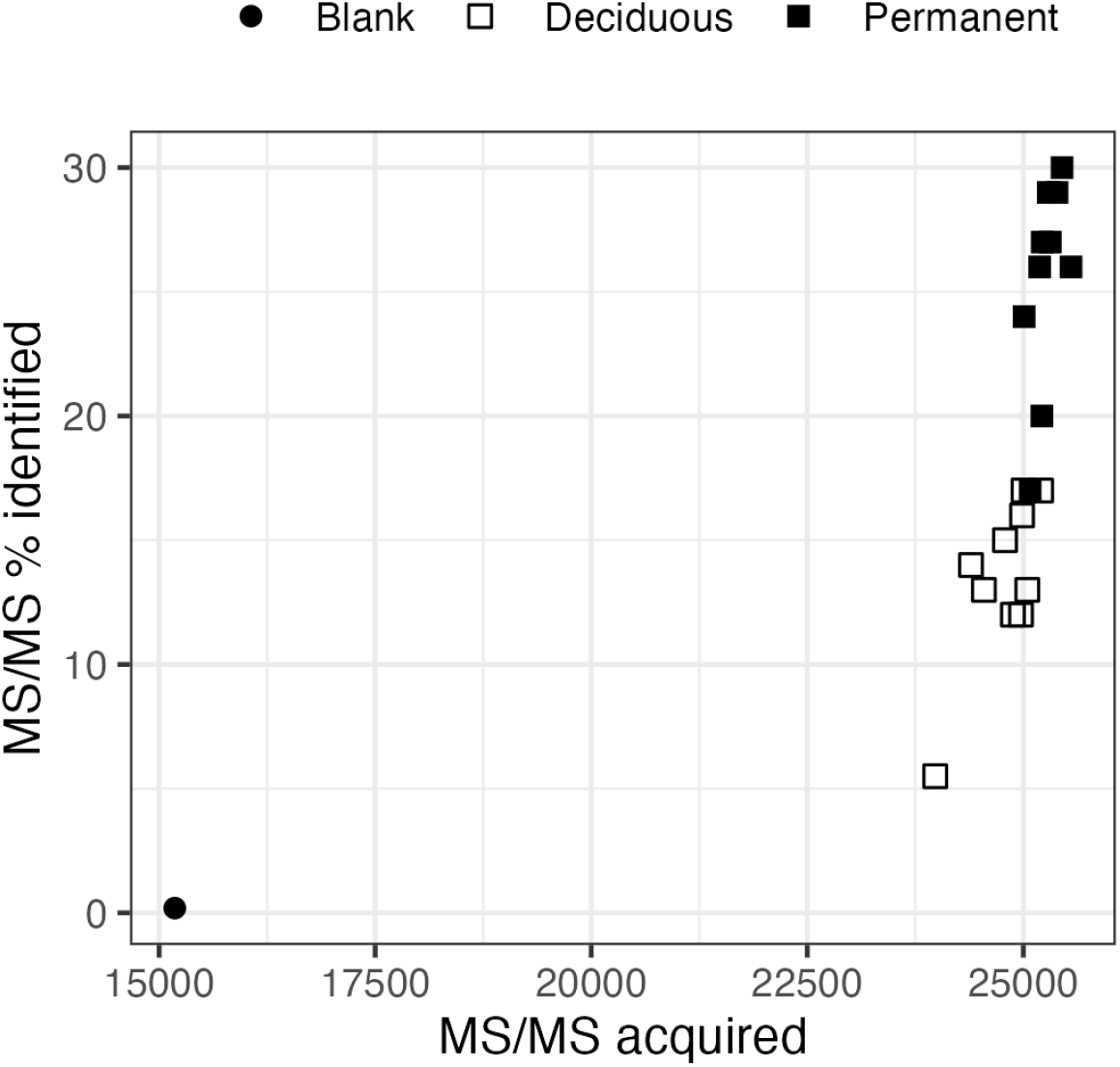
Quantitative comparison of MS/MS acquired and percentage of MS/MS identified between the laboratory blank (circle) and enamel extracts (deciduous teeth extracts in open squares and permanent teeth extracts in filled squares).

Twelve enamel proteins are identified in the 20 dental enamel extracts, where amelogenin X (AMELX), ameloblastin (AMBN), and enamelin (ENAM) are the most abundant. For AMELX and its two isoforms, the amino acid sequences are largely the same. Peptides found in all isoforms are in MaxQuant assigned to the most abundant isoform, in this case isoform 3 for both deciduous and permanent teeth and for both sexes. The same is observed for AMELY, where isoform 1 is the most abundant. The three least abundant proteins are matrix metalloproteinase 20 (MMP20), kallikrein 4 (KLK4), and odontogenic ameloblast-associated protein (ODAM), all of which are observed in higher peptide abundances in deciduous teeth (Table 1; SI Figure 1). We note that for the other core enamel proteins, protein abundance measured through protein intensity appears relatively similar between deciduous and permanent teeth. In contrast, peptide counts vary, with higher counts observed for AMTN in deciduous teeth compared to permanent teeth. The reverse is observed for AMBN, AMELX, AMELY, ENAM and COL17A1 (SI Figure 2). From the PEAKS search against the full human proteome, we confidently observe a further 92 proteins in the dataset with a minimum of 2 peptides in at least one specimen (SI Figure 3). Within the archaeological permanent dental enamel proteomes we observe proteins related to the immune system (e.g. various Immunoglobulins lambda constants (IgLs) and SERPINC1) which are not observed consistently in the modern deciduous dental enamel proteomes. Whether these proteins are endogenous or exogenous to the dental enamel or originate from underlying tissues or the oral cavity is currently unclear. Observations of these additional proteins in this study is in line with various additional modern and archaeological studies that also raise the possibility that the dental enamel proteome is larger than commonly assumed (Buonasera et al., 2024; Castiblanco et al., 2015; Gil-Bona & Bidlack, 2020; Green et al., 2019; Jágr et al., 2019; Lima Leite et al., 2018; Rexhaj et al., 2023).

**Table 1:** Mean number of unique peptides, with one standard deviation, of each of the core enamel proteins and identified isoforms for male and female deciduous and permanent teeth. An asterisk (*) indicates instances where SD could not be calculated.

| Protein | Accession | Deciduous |  | Permanent |  |
| --- | --- | --- | --- | --- | --- |
|  |  | Female | Male | Female | Male |
| ALB | P02768 | 11.4 ± 6.3 | 6.6 ± 3.4 | 12.0 ± 9.9 | 19.2 ± 16.0 |
| AMBN | Q9NP70 | 7.6 ± 3.2 | 8.8 ± 2.2 | 20.0 ± 4.2 | 19.5 ± 6.7 |
| AMELX | Q99217 | 10.2 ± 7.0 | 8.8 ± 4.6 | 27.5 ± 6.2 | 25.2 ± 6.6 |
| AMELX-2 | Q99217-2 | 3.5 ± 0.7 | 0 ± 0 | 3.5 ± 2.1 | 2.5 ± 0.6 |
| AMELX-3 | Q99217-3 | 20.6 ± 12.0 | 20.2 ± 8.6 | 62.5 ± 2.9 | 46.3 ± 12.2 |
| AMELY-1 | Q99218-1 | 0 ± 0 | 6.0 ± 1.8 | 0 ± 0 | 11.3 ± 3.3 |
| AMTN | Q6UX39 | 6.5 ± 2.9 | 12.8 ± 3.7 | 7.0* | 4.0 ± 1.0 |
| COL17A1 | Q9UMD9 | 17.0 ± 9.4 | 20.8 ± 7.2 | 86.5 ± 30.3 | 61.0 ± 19.5 |
| ENAM | Q9NRM1 | 148.4 ± 71.4 | 194.0 ± 39.0 | 443.0 ± 54.5 | 414.8 ± 79.8 |
| KLK4 | Q9Y5K2 | 4.0 ± 1.7 | 6.6 ± 3.9 | 0 ± 0 | 0 ± 0 |
| MMP20 | O60882 | 6.4 ± 4.0 | 7.4 ± 3.6 | 4.2 ± 2.1 | 5.2 ± 2.0 |
| ODAM | A1E959 | $3.5 \pm 2.1$ | $4.0 \pm 1.9$ | $2.0 \pm 0.0$ | 2.0* |

The two amelogenin proteins are coded for by genes on the sex chromosomes, AMELX on the X chromosome and AMELY on the Y chromosome (Lacruz et al., 2017; Santos & Line, 2006; Sire et al., 2007) and can therefore be used to estimate genetic sex. We confirmed the genetic sex for 19 out of the twenty individuals (Figure 3). This was done by comparing peptide intensities from label free quantification (LFQ) for two peptides containing complete or partial sequence for “SIRPPY**P**S” from AMELX and “S**M**IRPPY**S**S” from AMELY. These two sequences are the main difference between AMELX and AMELY, with the insertion of methionine (M) and substitution of proline (P) to serine (S) in AMELY. Otherwise the sequences are largely the same, which explains the identification of non-unique AMELY peptides in most specimens (Figure 2) as they are observed in both AMELX and AMELY. Estimated genetic sex from all permanent tooth extracts matches those of sex estimation based on skeletal morphology, six male extracts and four females. For the deciduous teeth, extracts of all female samples match what was self-reported upon tooth donation. AMELY-specific peptides were identified in all deciduous male teeth, confirming male genetic sex. For one specimen, individual 3, there was no LFQ intensity acquired for the single AMELY peptide (“SYEVLPLKWYQ**SM**”) identified in the MaxQuant search, where the last two amino acids are the start of the informative site (Figure 3). Therefore, its genetic sex cannot be estimated with certainty based on LFQ intensities alone, although the peptide is confidently identified. When comparing the results for this extract to the PEAKS searches with both isoforms of AMELY we observe five peptides containing peptides with the “SMIRPP” sequence, further supporting a male genetic sex assignment for the specimen. As a result, we found confirming proteomic evidence for the reported genetic sex for each individual.

**Figure 3:**
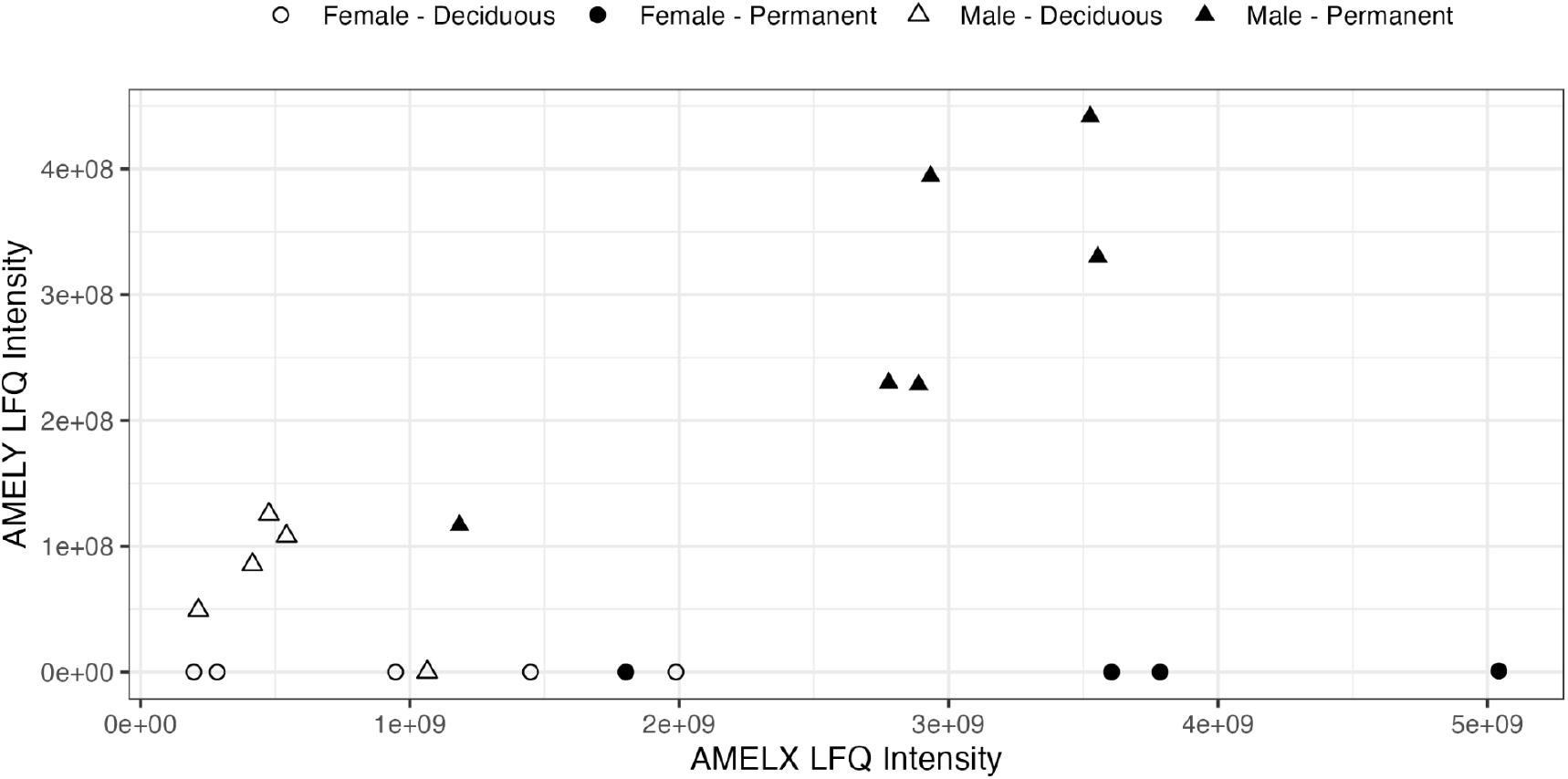
Estimation of genetic sex. The label-free quantification (LFQ) intensities represent a sum of intensities of peptides, containing complete or part of the sequence “SIRPPYPS” for AMELX and the sequence “SMIRPPYSS” for AMELY.

Due to the temporal and dentition differences between the two sample sets the following analyses are focused on the permanent dentition to highlight the preservation of *in vivo* influence of amelogenesis on the archaeological enamel proteome. Results from corresponding analysis for the deciduous dentition can be found in the supplementary material (SI Figures 4-7).

### 3.1. Effects of *in vivo* MMP20 and KLK4 digestion

The cleavage patterns induced by MMP20 and KLK4 during *in vivo* digestion for the most abundant enamel proteins, AMBN, AMELX, AMELY, and ENAM have been described in literature from experimental studies (Chun et al., 2010; Iwata et al., 2007; Nagano et al., 2009; Vetyskova et al., 2024; Yamakoshi et al., 2006, 2013). The survival of these cleavage patterns through time is largely unknown, with one study producing theoretical cleavage patterns based on Pleistocene enamel extracts (Welker et al., 2020).

#### 3.1.1. Ameloblastin

Cleavage of Ameloblastin (AMBN; Q9NP70) is a multifaceted process where not all cleavages are fully known (Chun et al., 2010). The initial hydrolysis events cleave AMBN into three protein products of different sizes, where we observe the N terminal 15 kDa cleavage product in the highest abundance (Figure 4) (Chun et al., 2010; Yamakoshi et al., 2001). We observe sequence coverage from position 27 to 127 that fall within the N-terminal 15 kDa cleavage product (the signal peptide covers positions 1 to 27 and is therefore not observed in the mature protein (Stakkestad et al., 2017)) (Figure 4). Additionally we observe peptides within the 23 kDa cleavage product, where we observe coverage for positions 223 to

**Figure 4:**
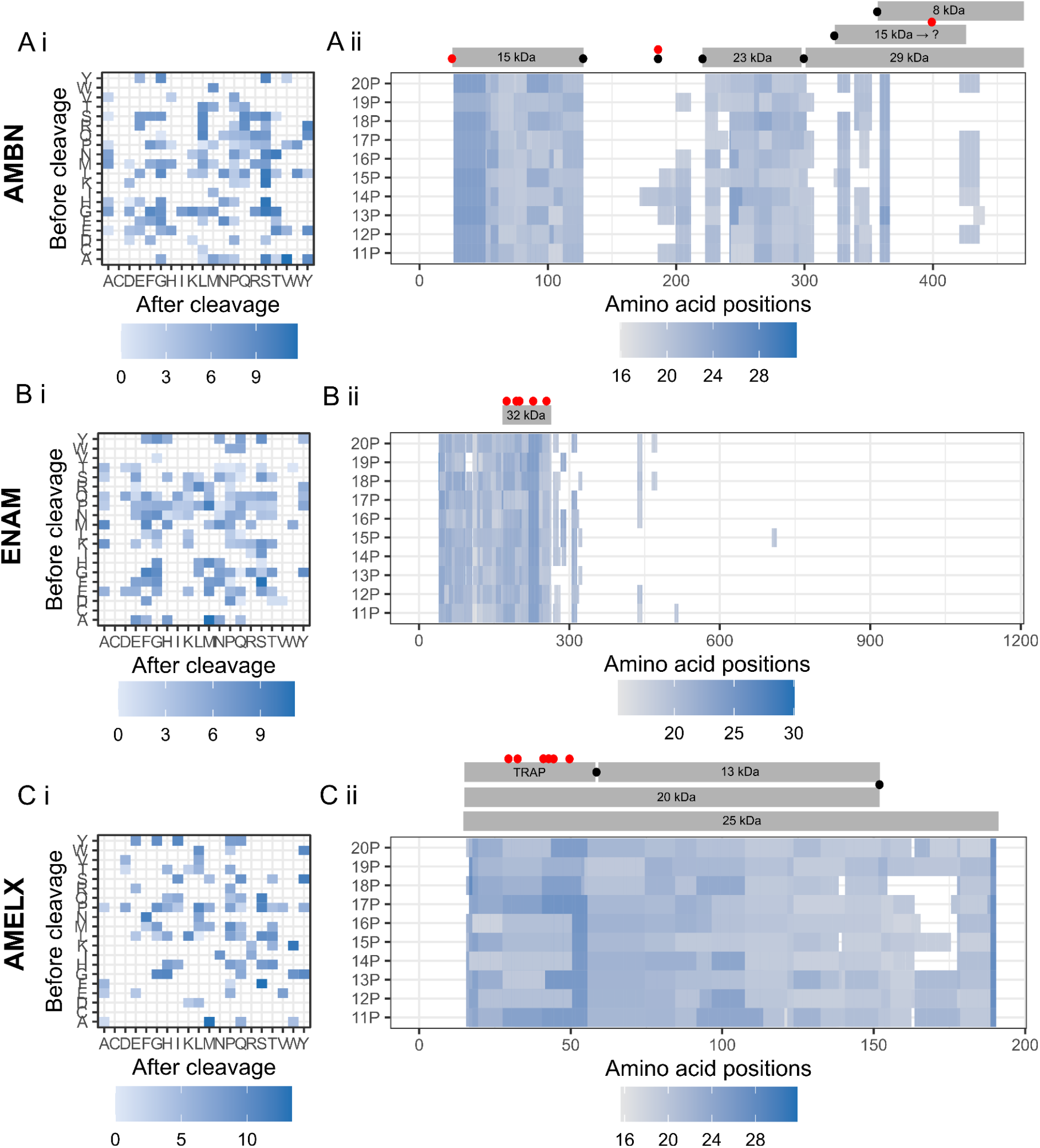
Influence of *in vivo* digestion of MMP20 and KLK4 for the ameloblast secreted proteins A) AMBN (Q9NP70), B) ENAM (Q9NRM1), C) AMELX (canonical sequence; Q99217). The influence is measured as i) observed cleavage sites per protein presented as a log2 fold change of observed PSM cleavage frequencies to all possible cleavages. As a result, higher values represent hydrolysis sites observed more frequently in the MS/MS data than randomly expected, ii) sequence coverage per protein shown as log2 of label free intensity (LFQ) of identified amino acid sequences. Annotations are based on porcine experimental data from AMBN (Chun et al., 2010); ENAM (Yamakoshi et al., 2006); AMELX (Nagano et al., 2009).

307. Between the MMP20 cleavages that give rise to the N-terminal 15 kDa and the 23 kDa cleavage products we observe fragmentary coverage. Finally, we observe coverage within the 29 kDa cleavage product with coverage of positions 322 to 366 and 421 to 440. This cleavage product undergoes additional cleavage by MMP20 and KLK4 and the coverage observed likely fall within the C terminal 15 kDa and 8 kDa cleavage products (Figure 4). These cleavage products have, however, not been fully characterised experimentally (Chun et al., 2010).

We observe overall cleavage to be the most frequent at the N terminal of serine (S), valine (V), tyrosine (Y), tryptophan (T), glycine (G), and at the C terminal of alanine (A), histidine (H), asparagine (N), glycine (G) and arginine (R). Cleavage was most often observed between serine and leucine (S:L) as well as glycine and leucine (G:L) as these amino acids occur next to each other ten and three times, respectively. Proportionally, cleavage between alanine and valine (A:V) was observed most often as these amino acids only occur next to each other once in the AMBN sequence.

#### 3.1.2. Enamelin

Enamelin (ENAM; Q9NRM1) is a rather large protein (1,142 amino acids) that contains a few cleavage products, mainly through MMP20 activity, the most stable one being the 32 kDa enamelin cleavage product (positions 136-241 in porcine ENAM) which is resistant to MMP20 cleavage and mainly undergoes cleavage by KLK4 (Yamakoshi et al., 2006). We observe coverage of the 32 kDa cleavage product, which is the most abundant region identified within enamelin in our data. Additionally, we observe coverage for the region between the signal peptide and the 32 kDa cleavage product (Figure 4). In the region C-terminal to the 32 kDa cleavage product we observe only sporadic sequence coverage. We observe overall cleavage to be the most frequent on the N terminal of methionine (M), serine (S), and phenylalanine (F) and on the C terminal of phenylalanine (F), alanine (A), glycine (G), and methionine (M). Cleavages are most often proportionally observed between alanine and methionine (A:M), and between phenylalanine and serine (F:S), which occur next to each other once each.

#### 3.1.3. Amelogenin

The *in vivo* digestion of amelogenin (both AMELX and AMELY) by MMP20 is known to produce a few cleavage products of varying size, based on porcine amelogenin. These cleavage products overlap each other for the most part (Nagano et al., 2009). We observe this as near complete sequence coverage for both amelogenin proteins, with highest coverage in the 6 kDa tyrosine rich amelogenin polypeptide (TRAP, amino acids 1-65; Figure 4). No major amelogenin cleavage products are known to be generated by KLK4. Which is, however, known to cleave mostly within the TRAP region (Nagano et al., 2009). Cleavages were overall observed most frequently at the N terminal of serine (S), methionine (M), tryptophan (W), and tyrosine (Y), as well as on the C terminal of phenylalanine (F), lysine (K), asparagine (N), and alanine (A). Cleavages are most observed between phenylalanine and serine (F:S), alanine and mathinone (A:M), lysine and tryptophan (K:W), which all occur only once each in the canonical AMELX sequence (Q99217) (Figure 4).

#### 3.1.4. Other enamel proteins

For the remaining enamel proteins identified within our dataset (AMTN (Q6UX39), ODAM (A1E959) and COL17A1 (Q9UMD9)) as well as the proteases themselves (MMP20 (O60882) and KLK4 (Q9Y5K2)) we identify rather fragmentary portions of the proteins (SI Figures 7-8). KLK4 was only observed in two permanent specimens with one peptide each and was therefore not studied further. Within ODAM, COL17A1, and KLK4 we observe higher abundance of cleavage on the N terminus of serine (S) compared to other amino acids (SI Figures 8-9).

### 3.2. Effects of FAM20C serine phosphorylation

The enamel proteins secreted by ameloblasts often undergo phosphorylation of the amino acid serine (S) induced by Family with sequence similarity member 20-C (FAM20C) (Dong et al., 2022; Ma et al., 2016; Tagliabracci et al., 2012). Sites of serine phosphorylation have been identified within AMBN, AMELX, AMELY, AMTN, and ENAM with the motifs of Serine - X - Glutamic acid (S-X-E) and Serine - X - phosphorylated-Serine (S-X-S(phos)), where X can be any amino acid (Brunati et al., 2000; Dong et al., 2022; Kawasaki & Weiss, 2003).

We observe a total of two to five sites of serine phosphorylation within the ameloblast secreted proteins (Figure 5). Two sites within AMBN not following the motif are serine at position 34, where the sequence is QQ**S**GT and where the threonine (T) is phosphorylated. Within AMELX, we observe one site at position 181 with a different motif, where the sequence is WP**S**TD, and where the serine (S) is phosphorylated. Whether these non-typical sites are from FAM20C phosphorylation, due to diagenesis, or due to another phosphorylation process is currently unknown.

**Figure 5:**
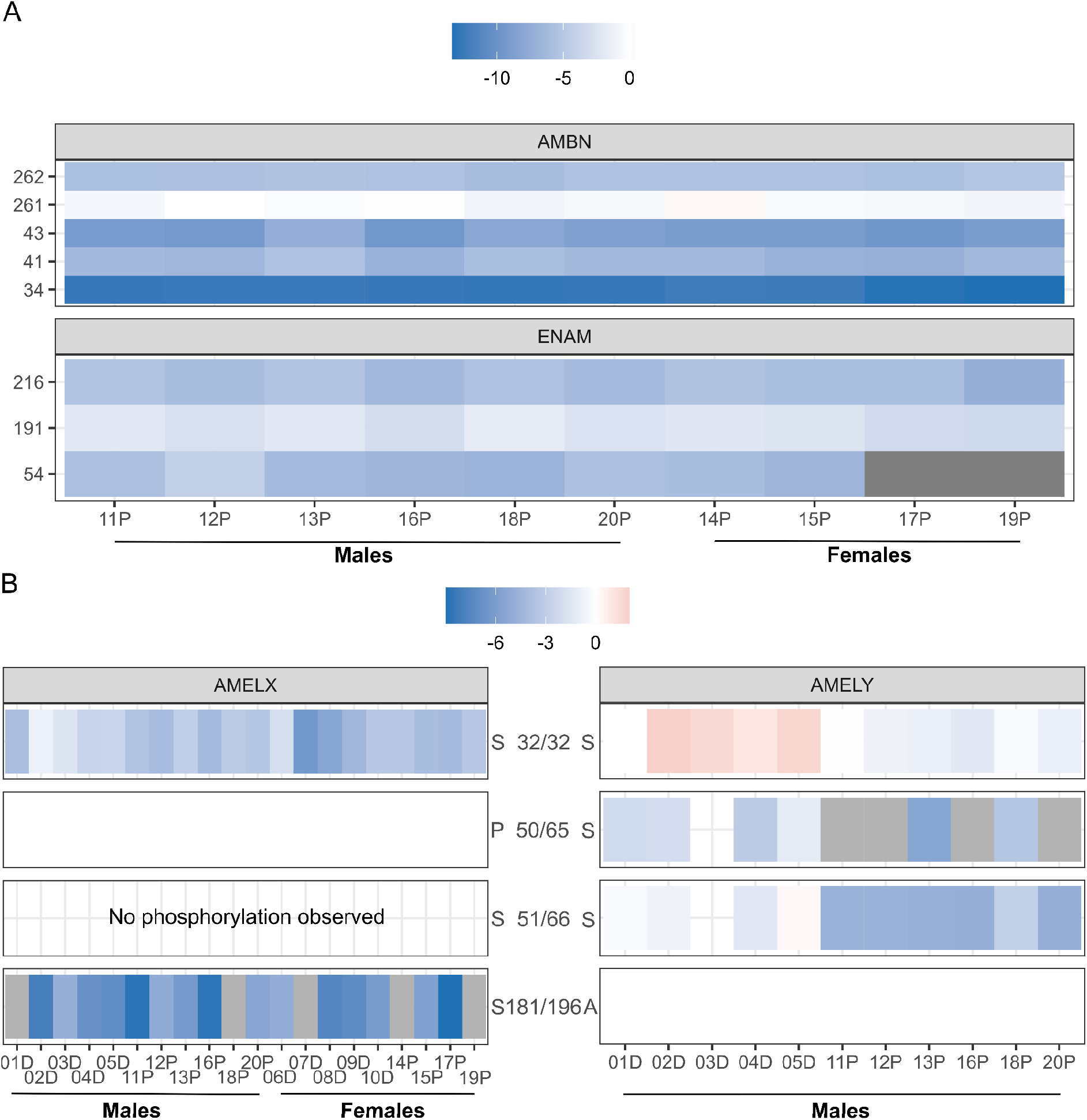
Phosphorylation sites observed within the enamel proteome for A) AMBN and ENAM, and B) AMELX (canonical sequence; Q99217) and AMELY (canonical sequence; Q99218-2). Phosphorylation is shown as log2 of the ratio of intensities from modified and unmodified peptides. Negative values indicate a higher amount of unmodified serines and positive values indicate a higher amount of phosphorylated serines. For AMELX and AMELY positions are based on positions within the respective canonical sequence. Grey bars indicate that the site is observed, but not with enough observations for quantification. Empty boxes indicate amino acids other than serine that do not undergo phosphorylation.

For these proteins we observe a similar ratio of modified to unmodified sites for both males and females for most sites with phosphorylation. The exception is position 66 in AMELY (“SM(ox)IRPPYS**S(phos)**”; Q99218). This position has previously been reported as phosphorylated only in male deciduous teeth, and has been proposed as a marker for male genetic sex, as the serine in the corresponding peptide for AMELX (“SIRPPYP**S**”) is not phosphorylated (Ziganshin et al., 2020). We observe phosphorylation at AMELY position 66 for both deciduous and permanent teeth, though the ratio of modified to unmodified sites is higher for deciduous teeth (Figure 5). For the corresponding serine in AMELX we observed no phosphorylation. This site can therefore be potentially used as a marker for male genetic sex for both deciduous and permanent teeth. Unfortunately, in our dataset we observe no peptides starting at the serine site with a phosphorylation - they are always non-phosphorylated. Furthermore, the serine phosphorylation on position 66 is only observed in peptides where position 66 is the C-terminus of the identified peptide. This might be due to MMP20 and/or KLK4 enzymatic activity (see above; (Welker et al., 2020)).

Additionally, the serine in position 32 in AMELY shows a higher ratio of phosphorylation for deciduous teeth compared to permanent teeth in males. Whether these differences are biological or induced by temporal differences between the dentition types is currently unknown, and could be explored in future studies of amelogenin phosphorylation rates in additional datasets.

## 4. Discussions

The enamel proteome, despite its small size, is widely used in palaeoproteomic research due to its incredible preservation over time, often outperforming the preservation of organic molecules in other mineralized tissues within the human body. Currently the enamel proteome is estimated to consist of nine to twelve proteins, with various other proteins reported to have been observed within the enamel proteome, often in archaeological material. Alongside the increase of enamel proteome use in phylogenetic research, its use in estimating genetic sex has expanded for both hominins and other mammals, using the two amelogenin proteins, AMELX and AMELY, coded for on the X and Y chromosome respectively (e.g. (Cappellini et al., 2019; Cintas-Peña et al., 2023; Koenig et al., 2024; Kotli et al., 2024; Lugli et al., 2019; Madupe et al., 2025; Parker et al., 2019; Stewart et al., 2016)). Different methods have been proposed for this estimation (see e.g. (Koenig et al., 2024; Parker et al., 2019)), however, there is no universal way of estimating genetic sex that applies to different enamel proteome extraction and mass spectrometry methods. Male genetic sex is determined by the presence of both AMELX and AMELY, while female genetic sex determination is more challenging to confidently assign, particularly as AMELY is ten times less abundant than AMELX (Santos & Line, 2006). This can therefore lead to false female genetic sex estimations when AMELY is not observed. We came across this challenge in the current study, when one tooth belonging to a modern individual with self-reported male genetic sex had a different genetic sex estimated based on peptide quantification, depending on which proteomic software was used to analyse the data. This highlights the need for a universal and statistically supported method for genetic sex estimation based on (palaeo)proteomic data.

For the ameloblast secreted proteins that undergo *in vivo* digestion by matrix metalloproteinase 20 (MMP20) and kallikrein 4 (KLK4), we observe the effects of the digestion in terms of cleavage patterns and which parts of the protein sequence are covered, as well as effects of serine phosphorylation by FAM20C, suggesting that the biology of enamel formation is still detectable in the proteome of archaeological remains. The *in vivo* digestion by MMP20 and KLK4 has mainly been studied experimentally on porcine enamel (Chun et al., 2010; Hu et al., 2002; Nagano et al., 2009; Yamakoshi et al., 2001, 2006, 2013). Additionally, it has been studied once in hominin palaeoproteomic data (Welker et al., 2020).

Ameloblastin (AMBN) is observed in three cleavage products, the N terminal 15 kDa cleavage products, the 23 kDa cleavage product along the middle of the protein sequence, as well as the C terminal 15 kDa cleavage product, with the first mentioned being most abundant. This matches largely to what has been reported in palaeoproteomic context (Welker et al., 2020). In enamelin (ENAM) is the 32 kDa cleavage product, as well as the region between the signal peptide and the 32 kDa cleavage product, the most abundant in our data. The remainder of the protein is rarely observed and only in a very fragmented state. That is in concordance with other experimental data as well as palaeoproteomic data (Welker et al., 2020; Yamakoshi et al., 2006), indicating that this abundant region preserves well through time and the remainder is either lost during amelogenesis or in too large fragments for a non-ezymatic protein extraction approach coupled to LC-MS/MS analysis. The amelogenin proteins (AMELX and AMELY) share the majority of their sequence. Together they are the most abundant proteins within our dental enamel extracts as well as having the highest sequence coverage, spanning all cleavage products of MMP20 digestion. This high sequence coverage has also been observed in hominin proteomes (Welker et al., 2020).

The effects of Family with sequence similarity member 20-C (FAM20C) serine phosphorylation in ameloblast secreted proteins are largely unknown in archaeological contexts. Here, we observe serine phosphorylations in AMBN, ENAM, AMELX, and AMELY that for the most part follow the “traditional” amino acid motifs of S-X-E and S-X-S(phos) (Brunati et al., 2000; Dong et al., 2022; Kawasaki & Weiss, 2003). These results are similar to those of hominin dental enamel proteomes (Welker et al., 2020) indicating biological origin rather than diagenetically-induced phosphorylation. We, however, also observe serine phosphorylations that do not follow the motifs, for example at position 34 of AMBN and position 181 in AMELX. Whether these serine phosphorylations are of biological or diagenetic origin should be explored in the future.

Additionally, we confirm a sex-specific serine phosphorylation in the amelogenin proteins. The most common region of the amelogenin proteins used to estimate genetic sex is “S-IRRPYPS” for AMELX (positions 44-51; Q99217) and corresponding “SM(ox)IRRPYSS” for AMELY (positions 58-66; Q99218) where the presence of the methionine insertion in AMELY is a determining factor. There is, however, corresponding serines at position 51 in AMELX and 66 in AMELY where we observe phosphorylation only in the male specimens. This has previously been described only in deciduous teeth (Ziganshin et al., 2020), We observe this phosphorylation in both deciduous and permanent male teeth.

Establishing how the *in vivo* biology of amelogenesis influences the archaeological dental enamel proteome, alongside diagenetic factors, is vital for palaeoproteomic research. Knowing the influence of MMP20 and KLK4 on the sequence coverage of proteins and which parts of the proteins are more stable and therefore more likely to preserve through time. Data analysis in palaeoproteomics is for the most part conducted through data dependent analysis (DDA) where protein sequences and post-translational modifications (PTMs) are selected prior to analysis. Including serine phosphorylation in that selection will allow for greater peptide identifications.

## 5. Conclusions

Here we explore the preservation of the *in vivo* digestion of MMP20 and KLK4, as well as FAM20C phosphorylation, during amelogenesis in archaeological material to gain further information on the dental enamel proteome. We demonstrate that the cleavage patterns of MMP20 and KLK4 digestion are preserved in the archaeological protein sequences of ameloblast secreted proteins. Where amelogenin proteins (AMELX and AMELY) show near complete sequence coverage over all cleavage products, while enamelin (ENAM) only shows sequence coverage in, and prior, to the 32 kDa cleavage product. This matches to prior porcine experimental data (Chun et al., 2010; Hu et al., 2002; Nagano et al., 2009; Yamakoshi et al., 2001, 2006, 2013), as well as hominin data (Welker et al., 2020). Additionally, we show serine phosphorylation patterns within the ameloblast secreted proteins in both permanent and deciduous dental enamel proteomes indicating that serine phosphorylation by FAM20C plays a vital role in amelogenesis and should be included as a post translational modification (PTM) in data analysis of dental enamel proteomes for increased peptide identification. We also identify a sex-specific serine phosphorylation marker in the AMELY sequence in both deciduous and permanent teeth. The results of this study of the dental enamel proteome opens up opportunities for improved datasets in the study of archaeological enamel proteomes, which in turn can aid in phylogenetic analysis.

## Acknowledgment

This research was supported by funding from the European Research Council (ERC) under the European Union’s Horizon 2020 research and innovation programme, grant agreement no. 948365 (PROSPER), awarded to F.W. F.W. and R.D.Á. are supported by the VILLUM FONDEN (no. 40747). M.M.T. and M.M.P. receive support from the Project PID2021-122355NB-C33 financed by MCIN/AEI/10.13039/501100011033/FEDER, UE.

M.M.P. has the support of the European Research Council within the European Union’s Horizon Europe (ERC-2021-ADG, Tied2-Teeth, project number 101054659). The Ratón Pérez project has been financed by the Spanish Foundation for Science and Technology (FECYT) of the Ministry of Science, Innovation, and Universities and Fundación “la Caixa”, Caixabank (FCT-20-15591). S.S. was funded by the Dutch Research Council (NWO) no. VI.Vidi.201.153). We thank Martin P. Defilet, senior management advisor for Heritage and Archaeology, and the city of Arnhem, the Netherlands, for access to skeletal remains. Mass spectrometry analysis, carried out at the Novo Nordisk Foundation Center for Protein Research, was funded in part by a donation from the Novo Nordisk Foundation (grant number NNF14CC0001).

