## Supplementary Material for "Archaeological preservation of amelogenesis pathways"

Ragnheiður Diljá Ásmundsdóttir[1], Gaudry Troché[1], Jesper V. Olsen[2], Marina Martínez de Pinillos[3], María Martinón-Torres[3], Sarah Schrader[4], Frido Welker[1]

1. Globe Institute, Faculty of Health and Medical Sciences, University of Copenhagen, Copenhagen, Denmark.
2. Novo Nordisk Foundation Center for Protein Research, Faculty of Health and Medical Sciences, University of Copenhagen, Copenhagen, Denmark.
3. Centro Nacional de Investigación sobre la Evolución Humana (CENIEH), Burgos, Spain
4. Laboratory of Human Osteoarchaeology, Faculty of Archaeology, Leiden University, Leiden, The Netherlands.

SI Table 1: Details of specimens used in current study. Biological sex of individuals donating deciduous teeth was self-reported. For the permanent teeth, biological sex was estimated based on skeletal morphology. Alphanumeric notation is used for tooth identification.

| **Individual** | **ID** | **Sample no.** | **Tooth** | **Dentition** | **Sex** | **Antiquity** | **Sample weight (mg)** |
|---|---|---|---|---|---|---|---|
| 1 | CENIEH-14/256-51 | 493 | LI1 | Deciduous | Male | Modern | <2 |
| 2 | CENIEH-14/412-204 | 494 | LI1 | Deciduous | Male | Modern | 11 |
| 3 | CENIEH-14/510-278 | 495 | LI1 | Deciduous | Male | Modern | 6.5 |
| 4 | CENIEH-14/475-292 | 496 | RI1 | Deciduous | Male | Modern | 2 |
| 5 | CENIEH-14/693-333 | 497 | LI1 | Deciduous | Male | Modern | 5 |
| 6 | CENIEH-14/246-16 | 498 | LI1 | Deciduous | Female | Modern | 5 |
| 7 | CENIEH-14/303-102 | 499 | LI1 | Deciduous | Female | Modern | 5 |
| 8 | CENIEH-14/492-283 | 500 | LI1 | Deciduous | Female | Modern | 5.5 |
| 9 | CENIEH-14/628-329 | 501 | LI1 | Deciduous | Female | Modern | 4 |
| 10 | CENIEH-1 | 502 | LI1 | Deciduous | Female | Modern | 5 |

| | | | | | | | |
|---|---|---|---|---|---|---|---|
| | 4/681-379 | | | | | | |
| 11 | V1621 | 503 | RI1 | Permanent | Male | Post-Medie val | 4 |
| 12 | V1428 | 504 | LI1 | Permanent | Male | Post-Medie val | <2 |
| 13 | V2190 | 505 | LI2 | Permanent | Male | Post-Medie val | 4 |
| 14 | V2193 | 506 | RI1 | Permanent | Female | Post-Medie val | 7 |
| 15 | V2324 | 507 | RI1 | Permanent | Female | Post-Medie val | 4 |
| 16 | V0868 | 508 | RI1 | Permanent | Male | Post-Medie val | 14 |
| 17 | V1506 | 509 | LI1 | Permanent | Female | Post-Medie val | 16 |
| 18 | V2141 | 510 | RI1 | Permanent | Male | Post-Medie val | 4 |
| 19 | V2351 | 511 | RI1 | Permanent | Female | Post-Medie val | <2 |
| 20 | V2406 | 512 | LI1 | Permanent | Male | Post-Medie val | <2 |

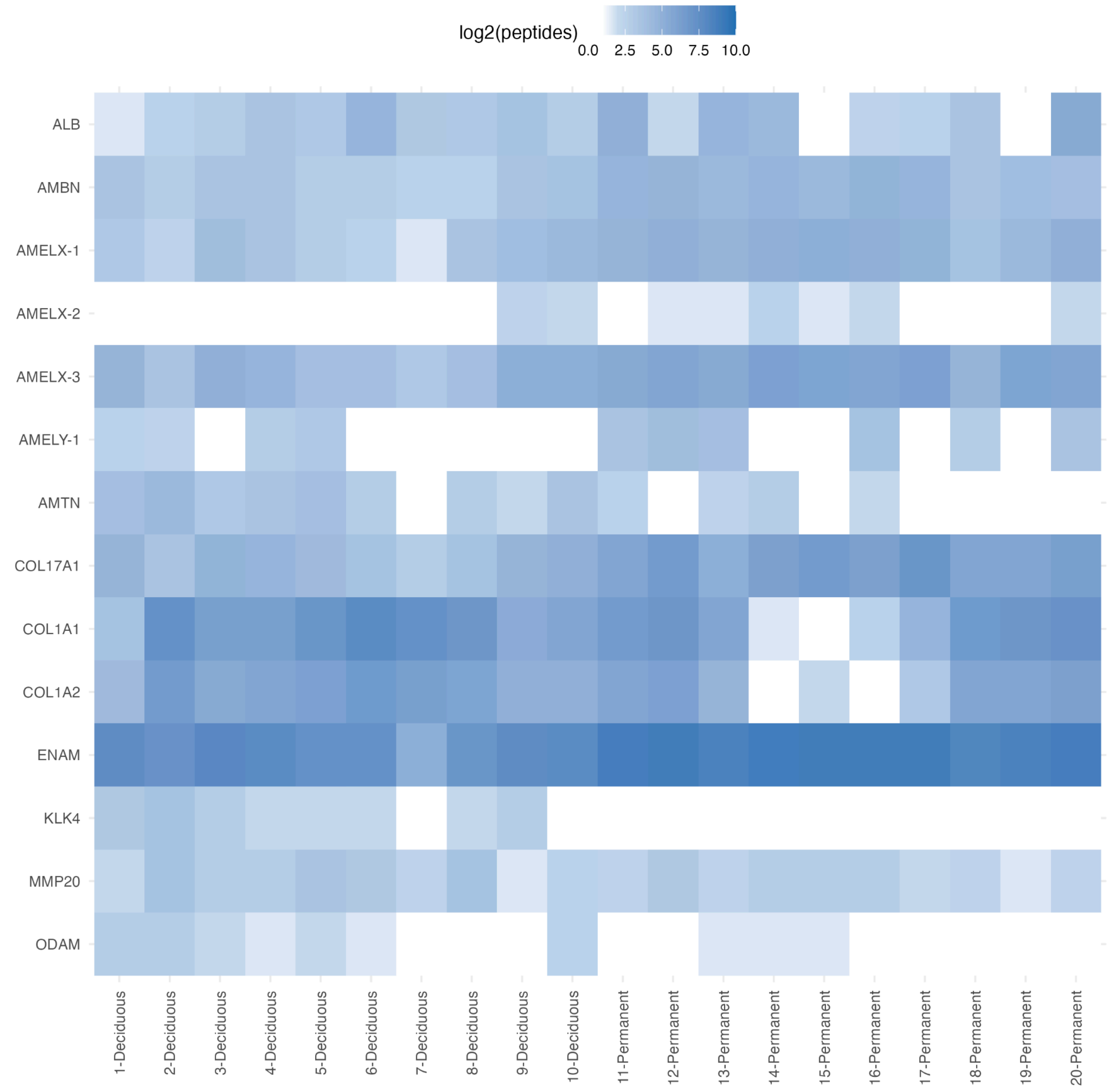


SI Figure 1: Proteome composition of deciduous and permanent dental enamel samples. Blank squares represent absence. A threshold of two unique peptides were required for protein identification per specimen.

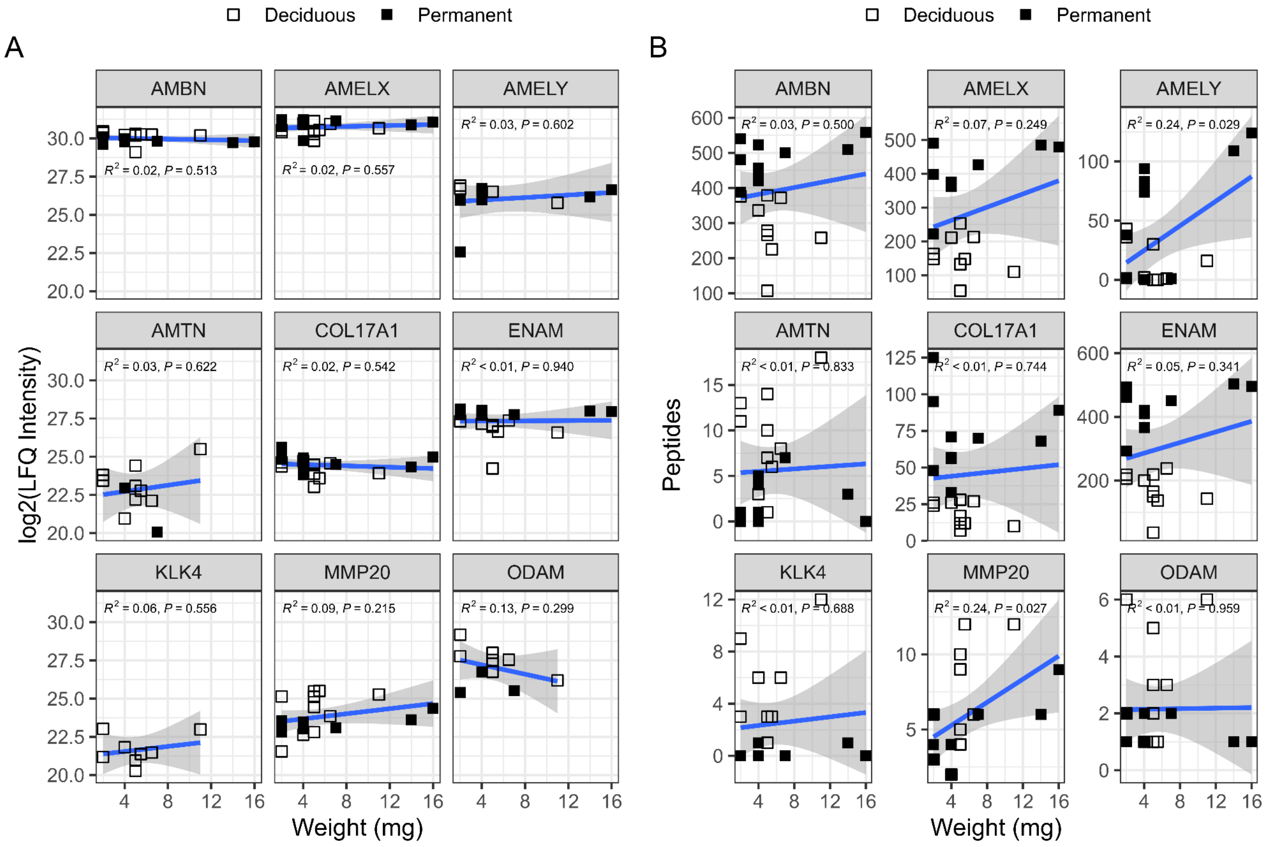


SI Figure 2: The effects of sample weight on core enamel proteome composition. A) Label-free quantification (LFQ) intensity. B) Number of peptides. Shaded area indicates 95% confidence interval.

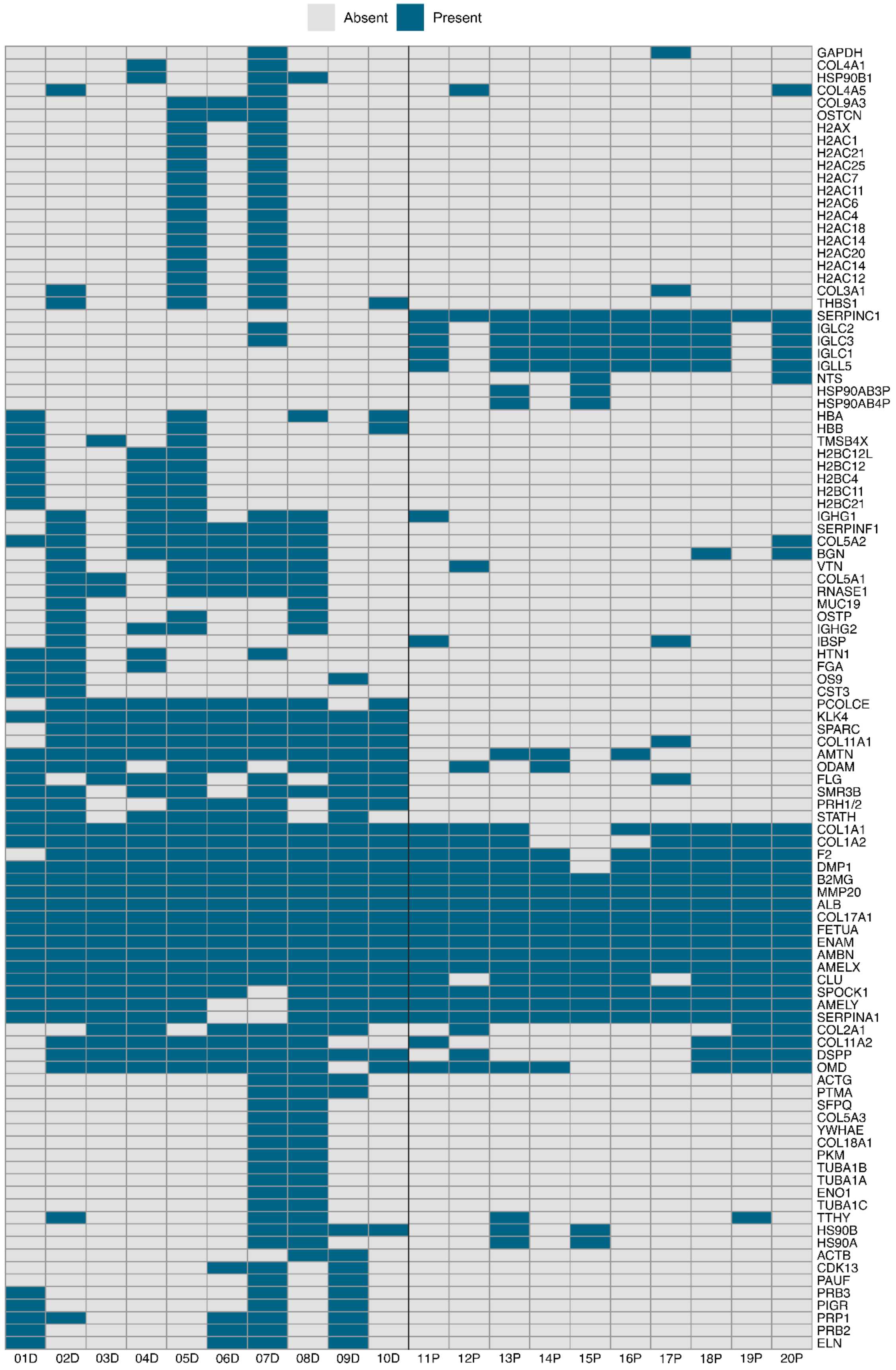


SI Figure 3: Presence and absence of identified proteins after the filtering process. Numbers on the x-axis indicate individuals (see SI Table 1). "D" indicates samples from deciduous teeth and “P” from permanent teeth.

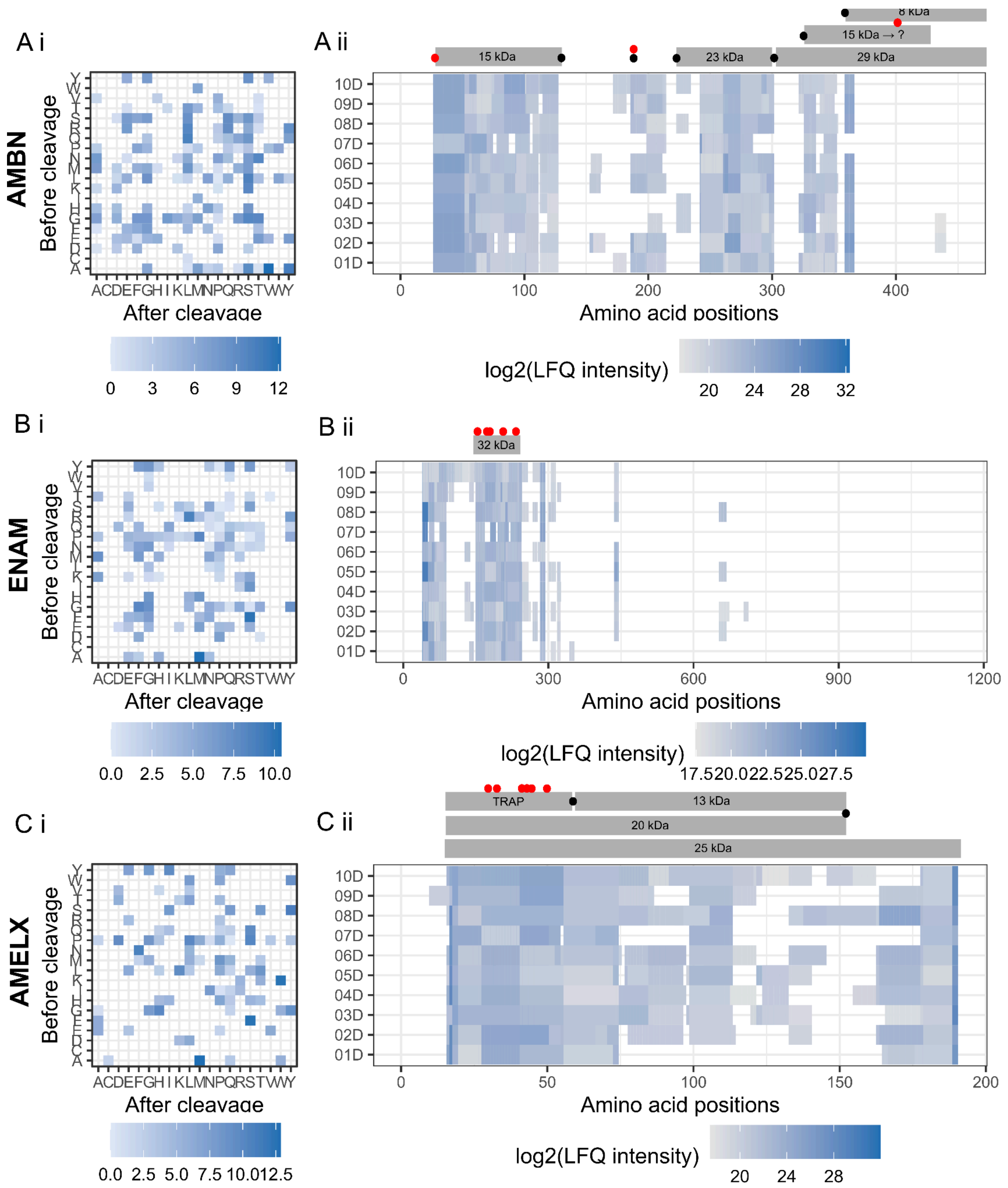


SI Figure 4: Influence of *in vivo* digestion of MMP20 and KLK4 for the ameloblast secreted proteins A) AMBN, B) ENAM, C) AMELX (canonical sequence; Q99217) in deciduous teeth. The influence is measured as i) observed cleavage sites per protein presented as a log2 fold change of observed PSM cleavage frequencies to all possible cleavages. As a result, higher values represent hydrolysis sites observed more frequently in the MS/MS data than randomly expected. ii) sequence coverage per protein shown as log2 of label free intensity (LFQ) of identified amino acid sequences. Annotations are based on porcine experimental data from AMBN (Chun et al. 2010); ENAM (Yamakoshi et al. 2006); AMELX (Nagano et al. 2009).

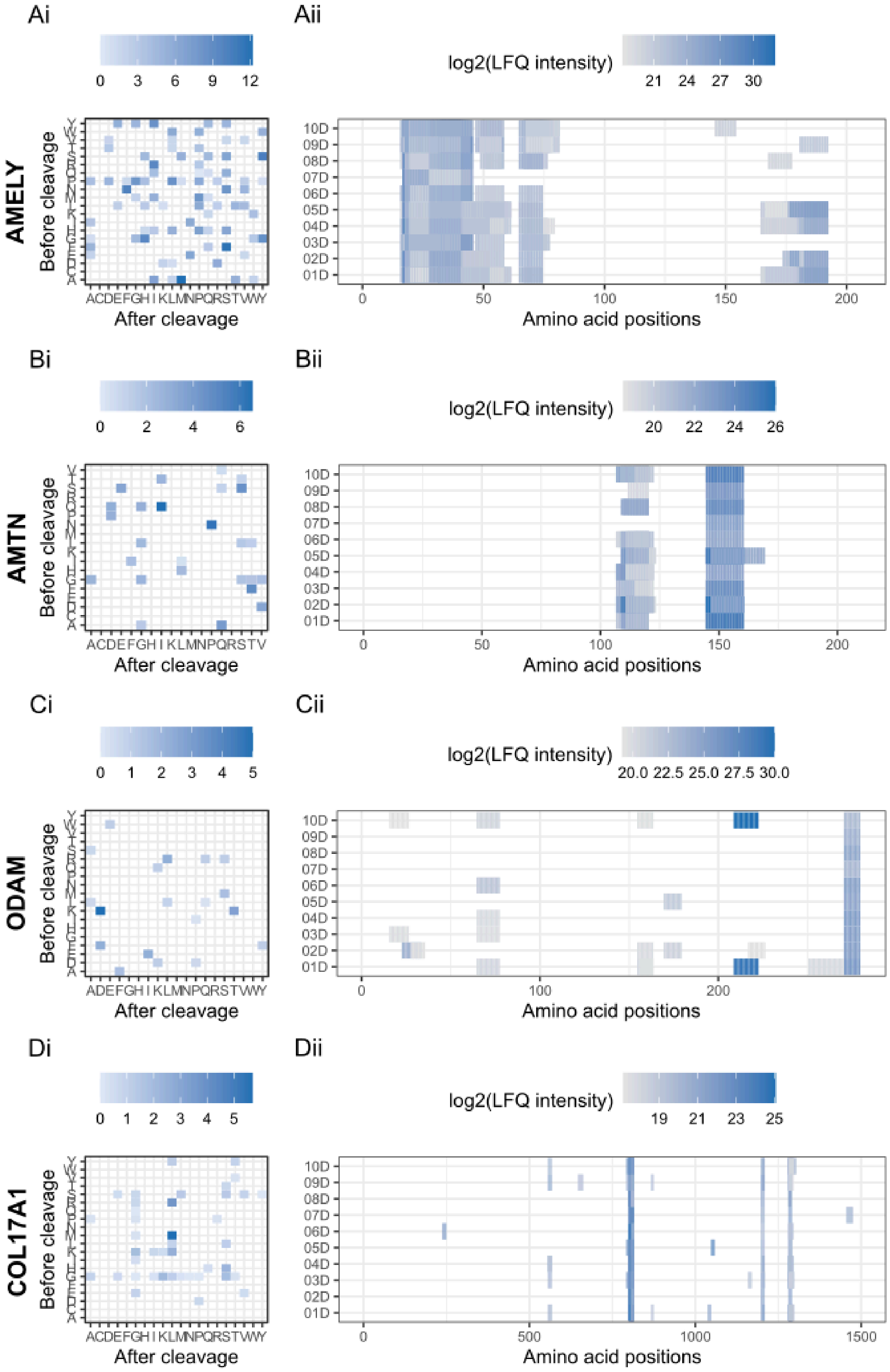


SI Figure 5: Influence of *in vivo* digestion of MMP20 and KLK4 for the proteins A) AMELY, B) AMTN, C) ODAM, and D) COL17A1 in deciduous teeth. The influence is measured as i) observed cleavage sites per protein presented as a log2 fold change of observed PSM cleavage frequencies to all possible cleavages. As a result, higher values represent hydrolysis sites observed more frequently in the MS/MS data than randomly expected. ii) sequence coverage per protein shown as log2 of label free intensity (LFQ) of identified amino acid sequences.

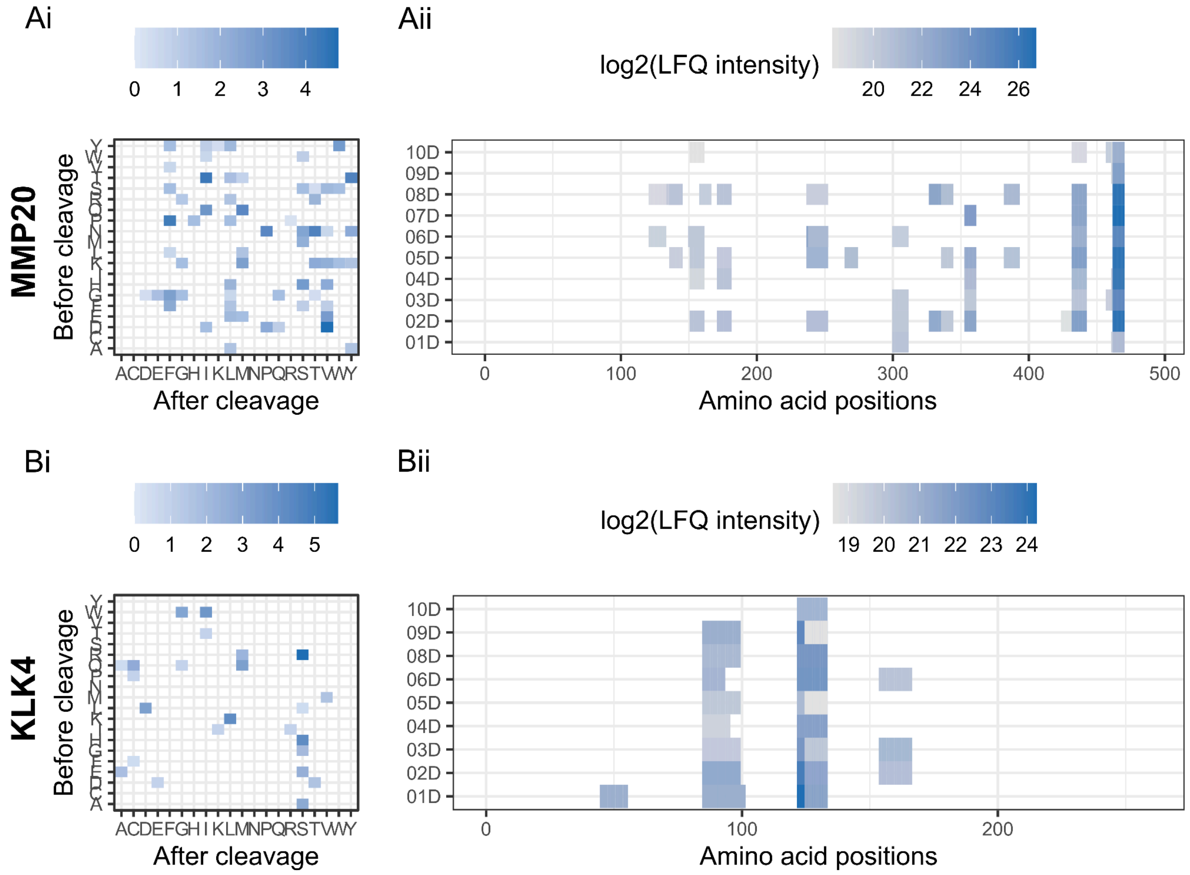


SI Figure 6: Preservation of A) MMP20 and B) KLK4 in deciduous dental enamel proteomes. i) observed cleavage sites per protein presented as a log2 fold change of observed PSM cleavage frequencies to all possible cleavages. As a result, higher values represent hydrolysis sites observed more frequently in the MS/MS data than randomly expected. ii) sequence coverage per protein shown as log2 of label free intensity (LFQ) of identified amino acid sequences.

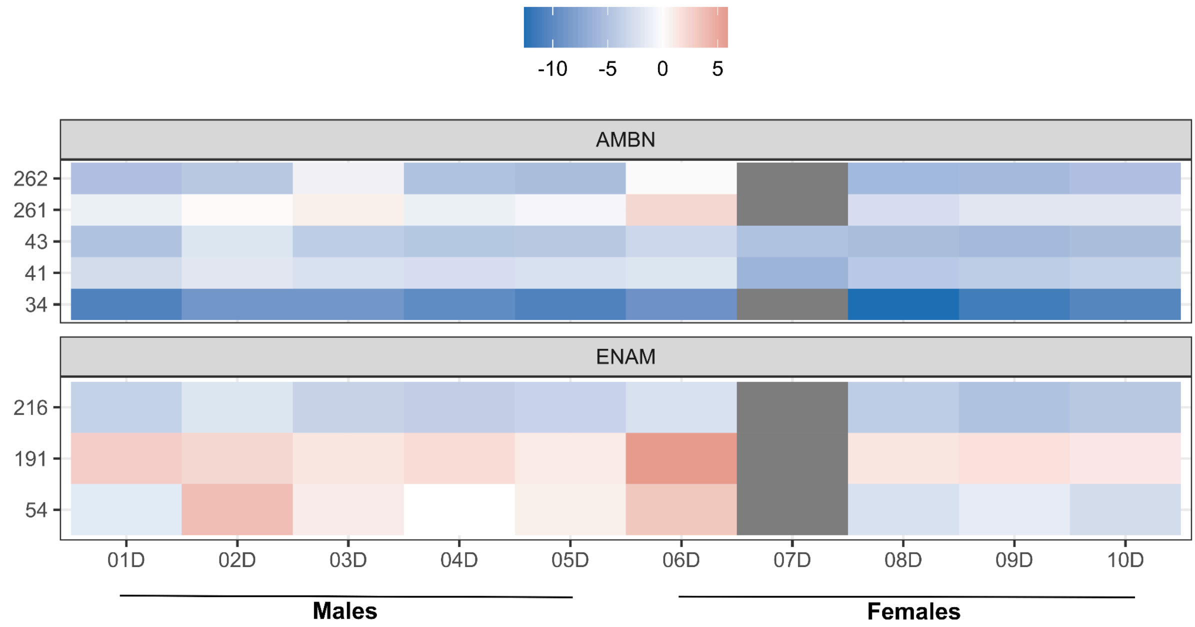

SI Figure 7: Phosphorylation sites observed within the deciduous dental enamel proteome for AMBN and ENAM. Phosphorylation is shown as log2 of the ratio of intensities from modified and unmodified peptides. Negative values indicate a higher amount of unmodified serines and positive values indicate a higher amount of phosphorylated serines.

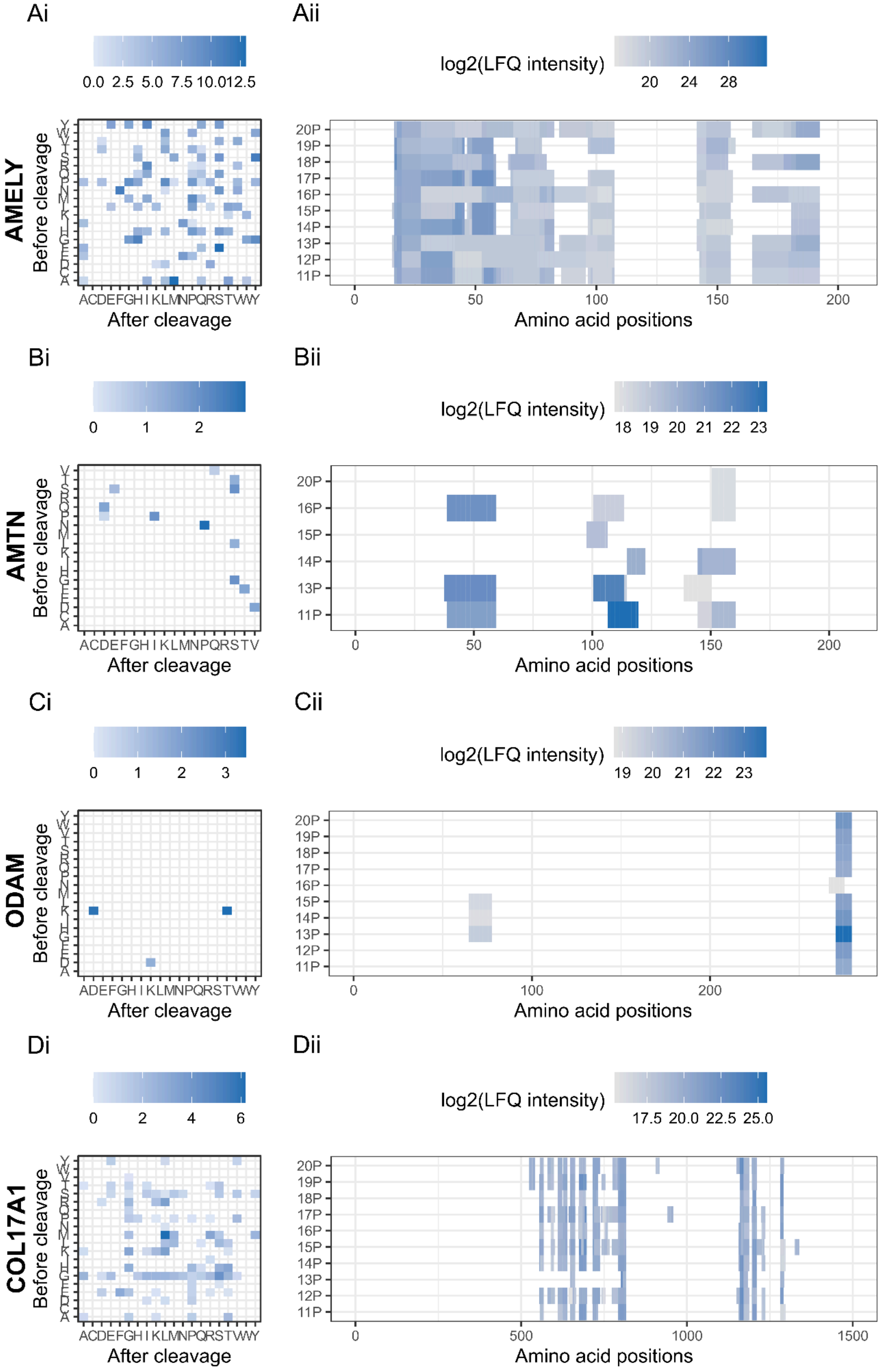

Ai
Aii
AMELY
Before cleavage
After cleavage
log2(LFQ intensity)
Amino acid positions
Bi
Bii
AMTN
Ci
Cii
ODAM
Di
Dii
COL17A1

SI Figure 8: Influence of *in vivo* digestion of MMP20 and KLK4 for the proteins A) AMELY, B) AMTN, C) ODAM, and D) COL17A1 in permanent teeth. The influence is measured as i) observed cleavage sites per protein presented as a log2 fold change of observed PSM cleavage frequencies to all possible cleavages. As a result, higher values represent hydrolysis sites observed more frequently in the MS/MS data than randomly expected. ii) sequence coverage per protein shown as log2 of label free intensity (LFQ) of identified amino acid sequences.

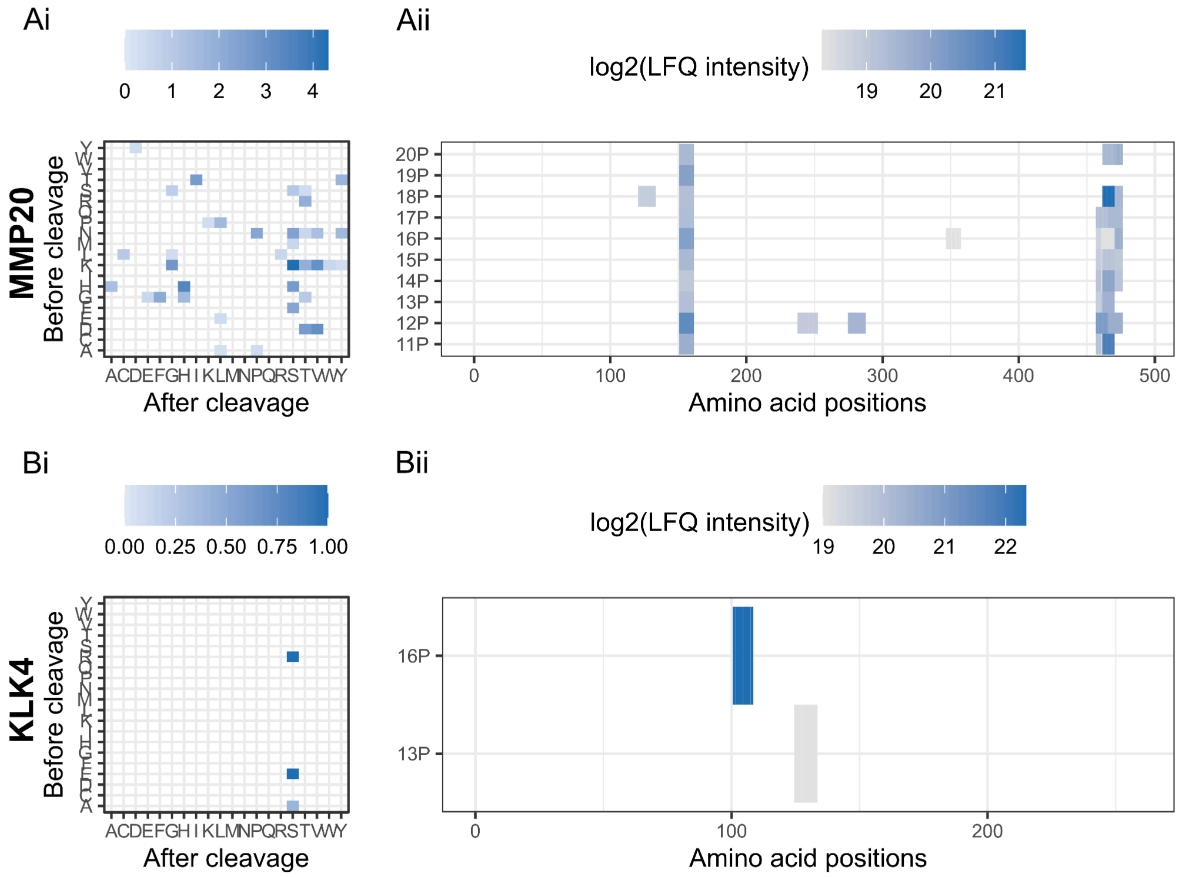


SI Figure 9: Preservation of A) MMP20 and B) KLK4 in permanent dental enamel proteomes. i) observed cleavage sites per protein presented as a log2 fold change of observed PSM cleavage frequencies to all possible cleavages. As a result, higher values represent hydrolysis sites observed more frequently in the MS/MS data than randomly expected. ii) sequence coverage per protein shown as log2 of label free intensity (LFQ) of identified amino acid sequences.
